# Google matrix analysis of bi-functional SIGNOR network of protein-protein interactions

**DOI:** 10.1101/750695

**Authors:** Klaus M. Frahm, Dima L. Shepelyansky

## Abstract

**Motivation:** Directed protein networks with only a few thousand of nodes are rather complex and do not allow to extract easily the effective influence of one protein to another taking into account all indirect pathways via the global network. Furthermore, the different types of activation and inhibition actions between proteins provide a considerable challenge in the frame work of network analysis. At the same time these protein interactions are of crucial importance and at the heart of cellular functioning.

**Results:** We develop the Google matrix analysis of the protein-protein network from the open public database SIGNOR. The developed approach takes into account the bi-functional activation or inhibition nature of interactions between each pair of proteins describing it in the frame work of Ising-spin matrix transitions. We also apply a recently developed linear response theory for the Google matrix which highlights a pathway of proteins whose PageRank probabilities are most sensitive with respect to two proteins selected for the analysis. This group of proteins is analyzed by the reduced Google matrix algorithm which allows to determine the effective interactions between them due to direct and indirect pathways in the global network. We show that the dominating activation or inhibition function of each protein can be characterized by its magnetization. The results of this Google matrix analysis are presented for three examples of selected pairs of proteins. The developed methods work rapidly and efficiently even for networks with several million of nodes and can be applied to various biological networks.

**Availability:** The Google matrix data and executive code of described algorithms are available at http://www.quantware.ups-tlse.fr/QWLIB/google4signornet/

## 1 Introduction

Protein-protein interactions (PPI) are at the heart of information processing and signaling in cellular functions. It is natural to present and analyze these PPI by presenting them as a directed network of actions between proteins (or network nodes). The simplest case of action is activation or inhibition so that such networks can be considered as bi-functional. The development of related academic databases of PPS networks with an open public access is a challenging task with various groups working in this direction (see e.g. Liberzon *et al*., 2011; Perfetto, L. *et al*., 2016; Kanehisa *et al*., 2017; Fabregat *et al*., 2018; Splender *et al*., 2018). A typical example is the SIGNOR directed network of PPI links for about 4000 proteins of mammals and 12000 bi-functional directed links as reported by Perfetto, L. *et al*., 2016.

On the scale of the past twenty years, modern society has created a variety of complex communication and social networks including the World Wide Web (WWW), Facebook, Twitter, Wikipedia. The size of these networks varies from a several millions for Wikipedia to billions and more for Facebook and WWW. The description of generic features of these complex networks can be found e.g. in Dorogortsev, 2010.

An important tool for the analysis of directed networks is the construction of the Google matrix of Markov transitions and related PageRank algorithm invented by Brin and Page in 1998 for ranking of all WWW sites (see Brin and Page, 1998; Langville and Meyer, 2006). This approach has been at the foundations of the Google search engine used world wide. A variety of applications of Google matrix analysis to various directed networks is described by Ermann *et al*., 2015.

Here we apply recently developed extensions of Google matrix analysis, which include the REduced GOogle MAtriX (REGOMAX) algorithm (Frahm *et al*., 2016) and the LInear Response algorithm for GOogle MAtriX (LIRGOMAX) (Frahm and Shepelyansky, 2019a), to the SIGNOR PPI network. The efficiency of these algorithms has been demonstrated for Wikipedia networks of politicians (Frahm *et al*., 2016) and world universities (Frahm and Shepelyansky, 2019a; Coquide *et al*., 2019a), and multi product world trade of UN COMTRADE database (Coquide *et al*., 2019b). Thus it is rather natural to apply these algorithms to PPI networks which have a typical size being significantly smaller than Wikipedia and WWW.

From a physical view-point the LIRGOMAX approach corresponds to a small probability pumping at a certain network node (or group of nodes) and absorbing probability at another specific node (or group of nodes). This algorithm allows first to determine the most sensitive group of nodes involved in this pumping-absorption process tracing a pathway connecting two selected proteins. In a second stage one can then apply the REGOMAX algorithm and obtain an effective reduced Google matrix, and in particular effective interactions, for the found subset of most sensitive nodes. These interactions are due to either direct or indirect pathways in the global huge network in which is embedded the selected relatively small subset of nodes.

The REGOMAX and LIRGOMAX algorithms originate from the scattering theory of nuclear and mesoscopic physics, field of quantum chaos and linear response theory of electron transport (Frahm *et al*., 2016; Frahm and Shepelyansky, 2019a).

We point out that the analysis of the SIGNOR PPI network already found biological applications reported by Sacco *et al*., 2016; Lun.X-K. *et al*., 2017; Kanhaiya *et al*., 2017; Dimitrakopoulos *et al*., 2018. The detailed review of various applications of the PPI signaling networks is given by Invergo and Beltrao, 2018. However, the Google matrix analysis has not been used in these studies.

The challenging feature of PPI networks is the bi-functionality of directed links which produce activation or inhibition actions. While in our previous analysis of SIGNOR network by Lages *et al*., 2018 this feature was ignored, here we apply the Ising-PageRank approach developed in Frahm and Shepelyansky, 2019b for opinion formation modeling. In this Ising-type approach the number of nodes in the PPI network is doubled, with a (+) or (*−*) attribute for each protein, and the links between doubled nodes are described by 2 *×* 2 matrices corresponding to activation or inhibition actions.

In this work we apply the LIRGOMAX and REGOMAX algorithm to the bi-functional PPI network of SIGNOR. We show that this approach allows to determine the effective sensitivity with direct and indirect interactions between a selected pair of proteins. As particular examples we will choose three protein pairs implicating the *Epidermal growth factor receptor (EGFR)* which is considered to play an important role in the context of lung cancer (see e.g. Bethune *et al*., 2010; Zamay *et al*., 2017).

The paper is constructed as follows: in Section 2 we describe the construction of Google matrix from links between proteins and related LIRGOMAX and REGOMAX algorithms, in Section 3 we characterize data sets and the Ising-PPI-network for bi-functional interactions between proteins, results are presented in Section 4 and the conclusion is given in Section 5. Supplementary Material provides additional matrix data and executable code for the described algorithms for the SIGNOR Ising-PPI-network.

## 2 Methods of Google matrix analysis

### 2.1 Google matrix construction

The Google matrix *G* of *N* nodes (proteins or proteins with (+)*/*(*−*) attribute) is constructed from the adjacency matrix *A*_*ij*_ with element 1 if node *j* points to node *i* and zero otherwise. The matrix *G* has the standard form *G*_*ij*_ = *αS*_*ij*_ + (1 − *α*)*/N* (see Brin and Page, 1998; Langville and Meyer, 2006; Ermann *et al*., 2015), where *S* is the matrix of Markov transitions with elements *S*_*ij*_ = *A*_*ij*_/*k*_*out*_(*j*) and 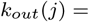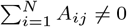 being the out-degree of node *j* (number of outgoing links); *S*_*ij*_ = 1*/N* if *j* has no outgoing links (dangling node). The parameter 0 < *α* < 1 is known as the damping factor with the usual value *α* = 0.85 (Langville and Meyer, 2006) which we use here. For the range 0.5 *≤ α ≤* 0.95 the results are not sensitive to *α* (Langville and Meyer, 2006; Ermann *et al*., 2015). A useful view on this *G* matrix is given by the concept of a random surfer, moving with probability *α* from one node to another via one of the available directed links or with a jump probability (1 − *α*) to any node.

The right PageRank eigenvector of *G* is the solution of the equation *GP* = *λP* for the leading unit eigenvalue *λ* = 1 (Langville and Meyer, 2006). The PageRank *P* (*j*) values represent positive probabilities to find a random surfer on a node *j* (∑_*j*_*P*(*j*) = 1). All nodes can be ordered by decreasing probability *P* numbered by PageRank index *K* = 1, 2, …*N* with a maximal probability at *K* = 1 and minimal at *K* = *N*. The numerical computation of *P*(*j*) is done efficiently with the PageRank iteration algorithm described by Langville and Meyer, 2006. The idea of this algorithm is simply to start with some initial, sum normalized, vector *P*^(0)^ of positive entries, e.g. being 1*/N* for simplicity, and then to iterate *P*^(*n*+1)^ = *G P*^(*n*)^ which typically converges after *n* = 150 − 200 iterations (for *α* = 0.85).

It is also useful to consider the original network with inverted direction of links. After inversion the Google matrix *G*\* is constructed via the same procedure with *G*\**P*\* = *P*\*. The matrix *G*\* has its own PageRank vector *P*\*(*j*) called CheiRank (Chepelianskii, 2010; Ermann *et al*., 2015). Its values give probabilities to find a random surfer of a given node and they can be again ordered in a decreasing order with CheiRank index *K*\* with highest *P*\* at *K*\* = 1 and smallest at *K*\* = *N*. On average, the high values of *P*(*P*\*) correspond to nodes with many ingoing (outgoing) links (Langville and Meyer, 2006; Ermann *et al*., 2015).

### 2.2 Reduced Google matrix (REGOMAX) algorithm

The REGOMAX algorithm is described in detail by Frahm *et al*., 2016; Lages *et al*., 2018; Coquide *et al*., 2019a. It allows to compute efficiently a “reduced Google matrix” *G*_R_ of size *N*_*r*_ × *N*_*r*_ that captures the full contributions of direct and indirect pathways appearing in the full Google matrix *G* between *N*_*r*_ nodes of interest selected from a huge global network with *N* ≫ *N*_*r*_ nodes. For these *N*_*r*_ nodes their PageRank probabilities are the same as for the global network with *N* nodes, up to a constant multiplicative factor taking into account that the sum of PageRank probabilities over *N*_*r*_ nodes is unity. The computation of *G*_R_ determines a decomposition of *G*_R_ into matrix components that clearly distinguish direct from indirect interactions: *G*_R_ = *G*_rr_ + *G*_pr_ + *G*_qr_ (Frahm *et al*., 2016). Here *G*_rr_ is given by the direct links between the selected *N*_*r*_ nodes in the global *G* matrix with *N* nodes. We note that *G*_pr_ is rather close to the matrix in which each column is approximately proportional to the PageRank vector *P*_r_, satisfying the condition that the PageRank probabilities of *G*_R_ are the same as for *G* (up to a constant multiplier due to normalization). Hence, in contrast to *G*_qr_, *G*_pr_ doesn’t give much new information about direct and indirect links between selected nodes.

The most interesting role is played by *G*_qr_, which takes into account all indirect links between selected nodes happening due to multiple pathways via the global network of nodes *N* (see Frahm *et al*., 2016). The matrix *G*_qr_ = *G*_qrd_ + *G*_qrnd_ has diagonal (*G*_qrd_) and non-diagonal (*G*_qrnd_) parts with *G*_qrnd_ describing indirect interactions between selected nodes. The exact formulas for all three components of *G*_R_ are given in Frahm *et al*., 2016. It is also useful to compute the weights *W*_R_, *W*_pr_, *W*_rr_, *W*_qr_ of *G*_R_ and its 3 matrix components *G*_pr_, *G*_rr_, *G*_qr_ given by the sum of all its elements divided by the matrix size *N*_*r*_. Due to the column sum normalization of *G*_R_ we obviously have *W*_R_ = *W*_rr_ + *W*_pr_ + *W*_qr_ = 1.

We note that the matrix elements of *G*qr may have negative values (only the full reduced matrix *G*_R_ should have positive elements; *G*_rr_ also has only positive matrix elements) but these negative values are found to be small for the Ising-PPI-networks and do not play a significant role. A similar situation for Wikipedia networks is discussed by Frahm *et al*., 2016; Frahm and Shepelyansky, 2019a.

### 2.3 LIRGOMAX algorithm

The detained description of the LIRGOMAX algorithm is given by Frahm and Shepelyansky, 2019a. It performs an infinitely w eak *ε*-probability injection (pumping) at one node (a protein or a protein with (+)/(−) attribute) and absorption at another node of interest. This process is described by the modified PageRank iteration *P*^(*n*+1)^ = *G F* (*ε, P* ^(*n*)^) where the vector valued function *F* (*ε, P*) has the components *P*(*i*) + *ε* for *i* being the index of the injection/pumping node, *P*(*j*) *− ε* for *j* being the index of the absorption node and simply *P*(*k*) for all other nodes *k*. In this way the vector *F* (*ε, P*) is also sum normalized if *P* is sum normalized and obviously *F* (0, *P*) = *P* is the identity operation. In Frahm and Shepelyansky, 2019a a more general version of *F*(*ε, P*) was considered with potentially different prefactors for the *ε* contributions, injection/absorption at possibly more than two nodes and an additional renormalization factor to restore the sum normalization (which is automatic in the simple version). However, for the applications in this work the above given simple version of *F* (*ε, P*) is sufficient.

In principle one can solve iteratively the above modified PageRank iteration formula which converges at the same rate as the usual PageRank iteration algorithm and provides a modified *ε*-depending PageRank *P* (*ε*). Then one can compute the linear response vector *P*_1_ = *dP* (*ε*)*/dε*|_*ε*=0_ =lim_*ε*→0_[*P*(*ε*) − *P*_0_]*/ε* where *P*_0_ is the PageRank obtained for *ε* = 0. However the naive direct evaluation of this limit is numerically not stable in the limit *ε* → 0. Fortunately as shown by Frahm and Shepelyansky, 2019a it is possible to compute *P*_1_ directly in an accurate and efficient way by solving the inhomogeneous PageRank equation

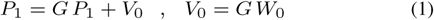

where the vector *W*_0_ has only two non-zero components for the two particular injection or absorption nodes *W*_0_(*i*) = 1 or *W*_0_(*j*) = −1 respectively. Therefore a more explicit expression for the vector *V*_0_ appearing in (1) is *V*_0_(*k*) = *G*_*ki*_ − *G*_*kj*_ (for all nodes *k*). We mention that the three vectors *P*_1_, *V*_0_ and *W*_0_ are orthogonal to the vector *E*^*T*^ = (1, …, 1) composed of unit entries, i.e. ∑_*k*_*P*_1_(*k*) = ∑_*k*_*W*_0_(*k*) = ∑_*k*_*V*_0_(*k*) = 0. Furthermore, all of these vectors, especially *P*_1_ have *real* positive or negative entries (note that in general eigenvectors of a non-symmetric real matrix may be complex).

A formal solution of the inhomogeneous PageRank equation is: 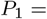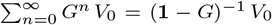 which is well defined since *V*_0_, when expanded in the basis of (generalized) eigenvectors of *G*, does NOT have a contribution of *P*_0_ (the only eigenvector of *G* with eigenvalue 1) such that the singularity of the matrix inverse does not constitute a problem. Of course numerically, we compute *P*_1_ in a different way, as described by Frahm and Shepelyansky, 2019a one can iterate the equation 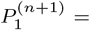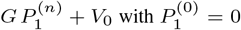 which converges with the same rate as the usual PageRank iteration.

In a similar as the PageRank *P*_0_ is characterized by the index *K* we introduce the index *K*_*L*_ by ordering *|P*_1_*|* such that *K*_*L*_ = 1 corresponds to the node with largest value of |*P*_1_| and *K*_*L*_ = *N* to the node with smallest value of |*P*_1_|. Once *P*_1_ is computed for the pair of chosen injection/absorption nodes we determine the 20 top nodes with strongest negative values of *P*_1_ and further 20 top nodes with strongest positive values of *P*_1_ which constitute a subset of 40 nodes which are the most significant nodes participating in the pathway between the pumping node *i* and absorbing node *j*. We also require that these two particular nodes *i* and *j* belong to this subset. If this is not automatically the case we replace the node at total position 20 (position 20 for strongest negative values of *P*_1_) with the absorption node *j* and/or the node at total position 40 (position 20 for strongest positive values of *P*_1_) with the injection node *i*. This situation happens once for the absorption node of the third example below which has a very low ranking position *K*_*L*_ ≈ 2000 for |*P*_1_|.

In general from a physical/biological point of view we indeed expect that the two particular injection/absorption nodes belong automatically to the selected subset of most sensitive nodes. However, there is no simple or general mathematical argument for this.

Using this subset of top nodes in the *K*_*L*_ ranking we then apply the REGOMAX algorithm to compute the reduced Google matrix and its components and in particular we determine the effective direct and indirect interactions of this reduced network. The advantage of the application of LIRGOMAX at the initial stage is that it provides an automatic and more rigorous procedure to determine an interesting subset of protein nodes related to the pumping between nodes *i* and *j* instead of using an arbitrary heuristic choice for such a subset.

## 3 Data sets and Ising-PPI-network construction

We use the open public SIGNOR PPI network Perfetto, L. *et al*., 2016 (April 2019 release for human, mouse and rat). This network contains *N* = 4341 nodes (proteins) and *N*_*ℓ*_ = 12547 directed hyperlinks between nodes. Each protein (node) is described by their name and identifier.

A new interesting feature of this PPI directed network is that its hyperlinks have activation and inhibition actions. For some links the functionality is unclear and then they are considered to be neutral. This feature rises an interesting mathematical challenge for the Google matrix description of such bi-functional networks. To meet this challenge we use the Ising-PageRank approach developed by Frahm and Shepelyansky, 2019b for a model of opinion formation on social networks. In this approach each node is doubled getting two components marked by (+) and (−). The activation links point to the (+) components and inhibition links point to the (−) components. Such transitions between doubled nodes are described by 2 × 2 block matrices *σ*_+_ (*σ*_−_) matrices with entries 1 (0) in the first row and 0 (1) in the second row as for Ising spin-1/2 (see details described in Supplementary Material). A neutral transition is described by 2 × 2 matrix *σ*_0_ with all elements being 1/2. Thus for this Ising-network (doubled-size network) we have doubled number of node *N* = 8682 and the total number of hyperlinks being *N*_*ℓ*_ = 27266; among them there are *N*_*act*_ = 14944 activation links, *N*_*inh*_ = 7978 inhibition links and *N*_*neut*_ = 4344 neutral links (*N*_*R*_ = *N*_*act*_ + *N*_*inh*_ + *N*_*neut*_). From this weighted Ising-PPI-network with *N*_*ℓ*_ = 27266 nodes we construct the Google matrix following the standard rules described by Langville and Meyer, 2006; Ermann *et al*., 2015 and also given above.

Below we apply the Google matrix analysis taking into account the bi-functionality PPI and illustrate the efficiency of the LIRGOMAX and REGOMAX algorithms for the SIGNOR Ising-PPI-network.

The details of Ising-PPI-network construction, its main statistical properties and an executable code for the described algorithms are provided in the Supplementary Material and in Frahm and Shepelyansky, 2019c. Below we discuss the results obtained with the LIRGOMAX and REGOMAX algorithms for three examples of specific pathways between two specific proteins.

## 4 Results

Here we present results obtained with LIRGOMAX and REGOMAX algorithms for pathways between several pairs of selected proteins.

### 4.1 Case of pathway EGFR - JAK2 proteins

As a first example we choose the node *EGFR P00533 (+)* for injection (pumping) and *JAK2 O60674 (-)* for absorption. It is known that mutations affecting the protein EGFR expression or activity could result in lung cancer (see e.g. Bethune *et al*., 2010; Zamay *et al*., 2017). This protein interacts with the protein JAK2 whose mutations have been implicated in various types of cancer. We argue that the injection (pumping) at *EGFR P00533 (+)* and absorption at *JAK2 O60674 (-)* should involve certain variations of the PageRank probability, represented by *P*_1_, showing interactions between various proteins actively participating in the pathway from *EGFR P00533 (+)* to *JAK2 O60674 (-)*. The pumping process can be viewed as a result of disease development and absorption as a certain mutation of this disease into another one.

The global PageRank indices of these two nodes are *K* = 90 (PageRank probability *P* (90) = 0.0009633) for *EGFR P00533 (+)* and *K* = 470 (PageRank probability *P* (470) = 0.0003444) for *JAK2 O60674 (-)*. As described above in the LIRGOMAX computations we choose the vector in *V*_0_ which appears in the inhomogeneous PageRank equation (1) as *V*_0_ = *G W*_0_ with *W*_0_(*K* = 90) = +1, *W*_0_(*K* = 470) = −1 and *W*_0_(*K*) = 0 for all other values of the Kindex *K*. We remind that both *W*_0_ and *V*_0_ are orthogonal to the left leading eigenvector *E*^*T*^ = (1, …, 1) of *G* according to the general description of the LIRGOMAX algorithm given above and in Frahm and Shepelyansky, 2019a.

For comparison we let us note that the top 4 PageRank nodes are *K* = 1 (*P* (1) = 0.003041) for *CASP3 P42574 (+)*, *K* = 2 (*P* (2) = 0.002821) for *NOTCH1 P46531 (+)*, *K* = 3 (*P* (3) = 0.002433) for *PIK3CD O00329 (-)*, *K* = 4 (*P* (4) = 0.002413) for *CTNNB1 P35222 (-)* (other values/data are available at Frahm and Shepelyansky, 2019c).

Similar to the two Wikipedia examples analyzed by Frahm and Shepelyansky, 2019a the LIRGOMAX algorithm selects the proteins mostly affected by injection/absorption process with 20 most positive (EGFR block) and 20 most negative (JAK2 block) values of *P*_1_ shown in Table 1. Here the pumped protein *EGFR P00533 (+)* is on the third position in its block of positive *P*_1_ values (*i* = 23) and with *K*_*L*_ = 11 (where *K*_*L*_ is the ranking index obtained by ordering the components of |*P*_1_|) while the protein with absorption *JAK2 O60674 (-)* has the second position in its block of negative *P*_1_ values (*i* = 2) with *K*_*L*_ = 4. Thus these two nodes are not at the first positions in their respective blocks but still they are placed at very high positions.

**Table 1.** Top 20 nodes of strongest negative values of *P*_1_ (index number *i* = 1*, …,* 20) and top 20 nodes of strongest positive values of *P*_1_ (index number *i* = 21, …, 40) with *P*_1_ being created as the linear response of the PageRank of the Ising-PPI-network with injection (or pumping) at *EGFR P00533* (+) and absorption at *JAK2 O60674 (-)*; *K*_*L*_ is the ranking index obtained by ordering |*P*_1_| and *K* is the usual PageRank index obtained by ordering the PageRank *P*_0_.

| $i$ | $K_L$ | $K$ | Node name |
| --- | --- | --- | --- |
| 1 | 1 | 30 | JAK2 O60674 (+) |
| 2 | 4 | 470 | JAK2 O60674 (-) |
| 3 | 5 | 554 | IFNGR2/INFR1 SIGNOR-C142 (+) |
| 4 | 6 | 354 | ARHGEF1 Q92888 (+) |
| 5 | 7 | 631 | APOA1 P02647 (+) |
| 6 | 8 | 956 | CSF2RA/CSF2RB SIGNOR-C212 (+) |
| 7 | 9 | 57 | STAT1 P42224 (+) |
| 8 | 10 | 204 | MAP3K5 Q99683 (-) |
| 9 | 12 | 1008 | STAT4 Q14765 (+) |
| 10 | 13 | 825 | CCR2 P41597 (+) |
| 11 | 14 | 2377 | PRMT5 O14744 (-) |
| 12 | 15 | 2378 | STAM Q92783 (+) |
| 13 | 16 | 1482 | EPOR P19235 (+) |
| 14 | 17 | 1117 | CSF2RA P15509 (+) |
| 15 | 18 | 959 | ITGAL P20701 (+) |
| 16 | 19 | 1968 | CTLA4 P16410 (-) |
| 17 | 20 | 2058 | STAP2 Q9UGK3 (+) |
| 18 | 21 | 2024 | ITGB2 P05107 (+) |
| 19 | 22 | 532 | EZH2 Q15910 (-) |
| 20 | 23 | 1196 | GTF2I P78347 (+) |
| 21 | 2 | 29 | GRB2 P62993 (+) |
| 22 | 3 | 172 | FES P07332 (+) |
| 23 | 11 | 90 | EGFR P00533 (+) |
| 24 | 27 | 3 | PIK3CD O00329 (-) |
| 25 | 30 | 126 | CBL P22681 (+) |
| 26 | 31 | 136 | EGFR P00533 (-) |
| 27 | 32 | 648 | EZR P15311 (+) |
| 28 | 36 | 38 | PTK2 Q05397 (+) |
| 29 | 37 | 456 | GAB1 Q13480 (+) |
| 30 | 39 | 424 | BCR P11274 (-) |
| 31 | 42 | 124 | PIK3R1 P27986 (+) |
| 32 | 43 | 26 | PLCG1 P19174 (+) |
| 33 | 44 | 58 | SHC1 P29353 (+) |
| 34 | 45 | 88 | ESR1 P03372 (+) |
| 35 | 46 | 2398 | VAV2 P52735 (+) |
| 36 | 47 | 746 | SHC3 Q92529 (+) |
| 37 | 48 | 291 | ERBB2 P04626 (+) |
| 38 | 49 | 888 | ERBB3 P21860 (+) |
| 39 | 51 | 1109 | NCK1 P16333 (+) |
| 40 | 52 | 1531 | CRK P46108 (-) |

The dependence of |*P*_1_| of the index *K*_*L*_ is shown in the top panel of Figure 1. The decay of |*P*_1_| is relatively slow for *K*_*L*_ ≤ 40 followed by a more rapid drop for *K*_*L*_ > 40. The bottom panel shows the dependence of positive (blue) and negative (red) values of *P*_1_ on *K*_*L*_. We note that the top absolute values |*P*_1_| for blue and red components have comparable values being of the order of |*P*_1_| ~ 0.1 for approximately *K*_*L*_ ≤ 40. However, in this range the number of positive (blue) values of *P*_1_ is significantly smaller compared to the number of negative (red) values of *P*_1_. This point can also be seen from the column of *K*_*L*_ values in Table 1. Another feature visible from Table 1 is that the number of proteins with negative component (−) is significantly smaller than those with a positive component (+) (5 for 1 ≤ *i* ≤ 20 and 4 for 21 ≤ *i* ≤ 40. We return to the properties of positive and negative components a bit later.

**Fig. 1.**
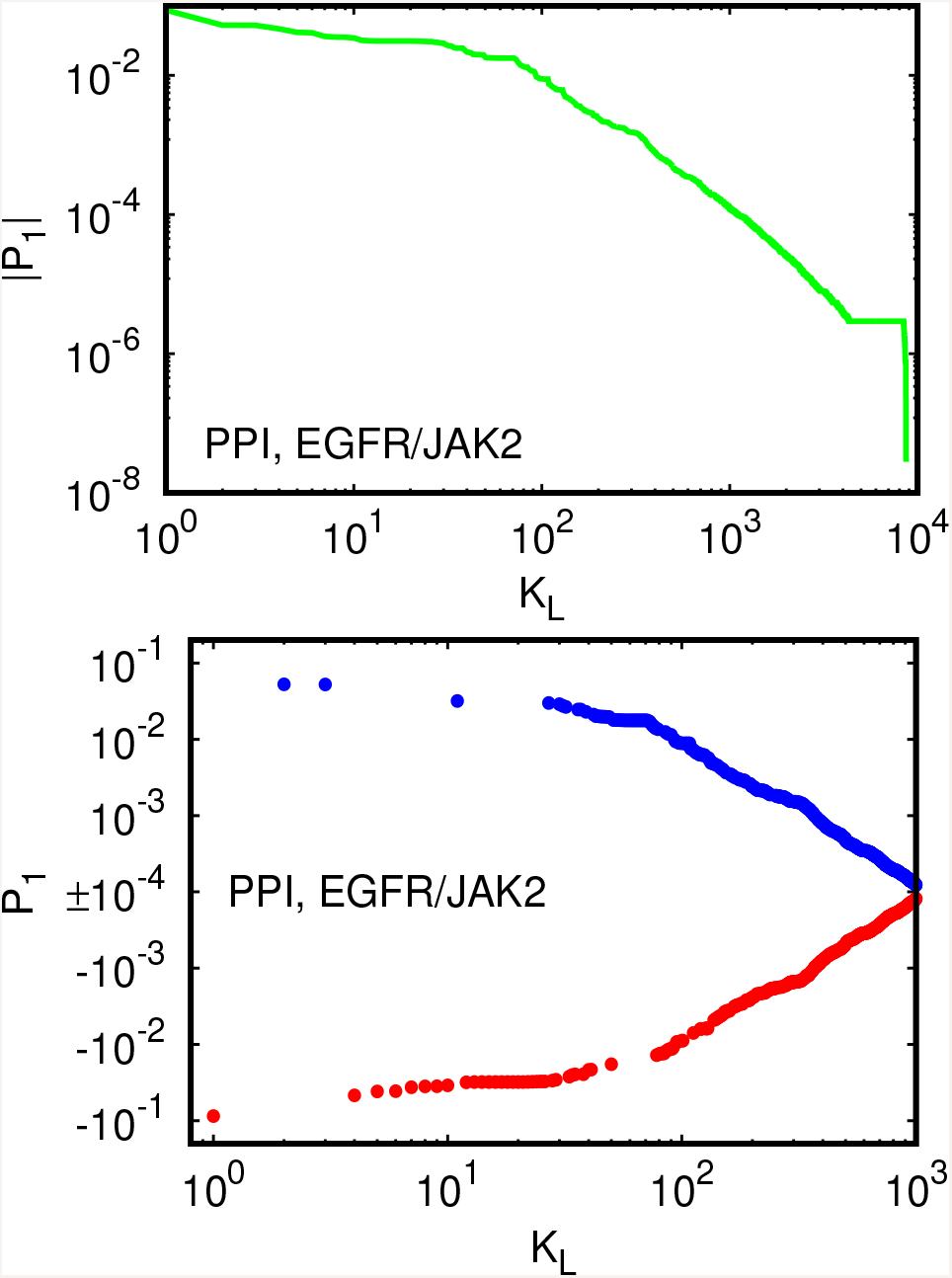
Linear response vector *P*_1_ of PageRank for the Ising-PPI-network with injection (or pumping) at *EGFR P00533 (+)* and absorption at *JAK2 O60674 (-)*. Here *K*_*L*_ is the ranking index obtained by ordering |*P*_1_| from maximal value at *K*_*L*_ = 1 down to minimal value. Top panel shows |*P*_1_| versus *K*_*L*_ in a double logarithmic representation for all *N* nodes. Bottom panel shows a zoom of *P*_1_ versus *K*_*L*_ for *K*_*L*_ ≤ 10^3^ in a double logarithmic representation with sign; blue data points correspond to *P*_1_ > 0 and red data points to *P*_1_ < 0.

After the selection of most significant 40 nodes of the pathway between the two injection/absorption proteins (see Table 1) we apply the REGOMAX algorithm which determines all matrix elements of Markov transitions between these 40 nodes including all direct and indirect pathways via the large global Ising-PPI-networks network with 8682 nodes.

The reduced Google matrix *G*_R_ and its three components *G*_pr_, *G*_rr_, *G*_qr_ are shown in Figure 2 for proteins of Table 1 (1 ≤ *i* ≤ 40). The weight of the component *G*_pr_ is *W*_pr_ = 0.761 being not so far from unity but this value is below *W*_pr_ ≈ 0.95 appearing usually in Wikipedia networks (Frahm *et al*., 2016; Frahm and Shepelyansky, 2019a). We attribute this to a significantly smaller number of links per node being *ℓ* = *N*_*ℓ*_/N ≈ 3.1 for the Ising-PPI-network while for the English Wikipedia network of 2017 we have *ℓ* ≈ 22.5 (Frahm and Shepelyansky, 2019a). Indeed, the weight *W*_rr_ = 0.220 of direct transitions of *G*_rr_ is significantly larger than the corresponding values for the Wikipedia case with *W*_rr_ ≈ 0.04. However, the weights *W*_qr_ = 0.019 are comparable for both reduced networks.

**Fig. 2.**
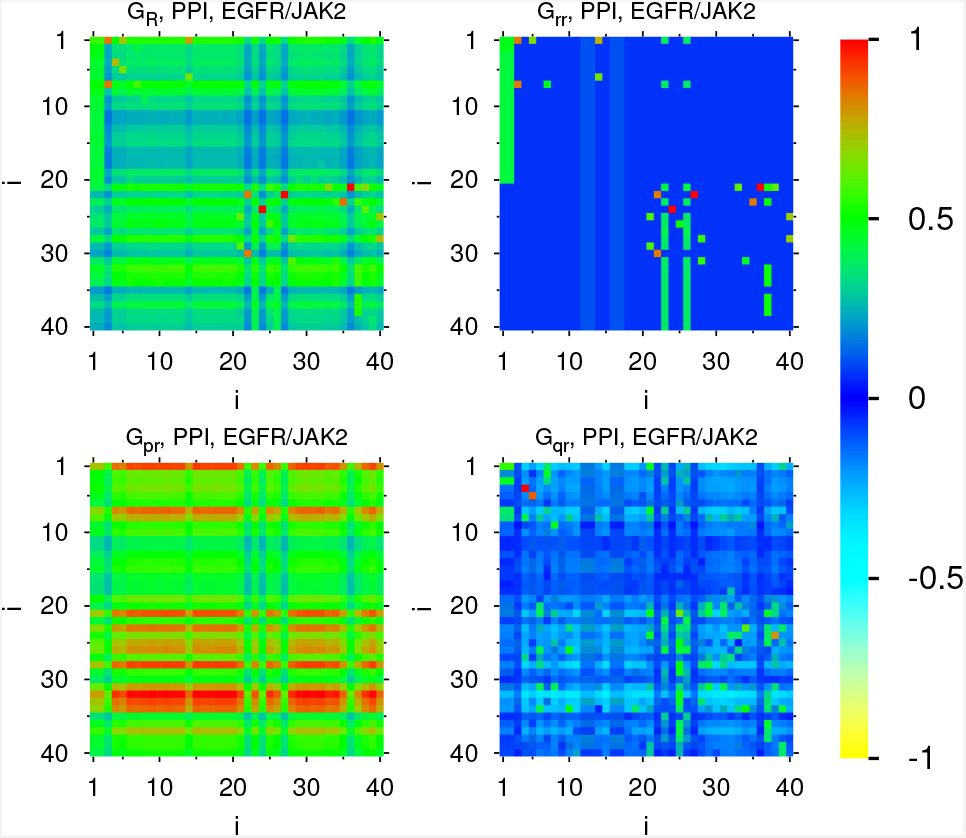
Reduced Google matrix components *G*_R_, *G*_pr_, *G*_rr_ and *G*_qr_ for Ising-PPI-network and the subgroup of nodes given in Table 1 corresponding to injection at *EGFR P00533 (+)* and absorption at *JAK2 O60674 (-)* (see text for explanations). The axis labels correspond to the index *i* used in Table 1. The relative weights of these components are *W*_pr_ = 0.761, *W*_rr_ = 0.220, and *W*_qr_ = 0.019. The values of the color bar correspond to sgn(*g*)(|*g*|/ max |*g*|)^1/4^ where *g* is the shown matrix element value. The exponent 1/4 amplifies small values of *g* for a better visibility.

The matrix structure of direct transitions *G*_rr_ has a clear two block structure with dominant transitions inside each block associated to EGFR and JAK2 with only 4 significant matrix elements from the EGFR to the JAK2 block. These matrix elements correspond to links from EGFR (*±*) to JAK2 (+) and STAT1(+) and have the same value *g* ≈ 0.0167 while all other matrix elements (of this EGFR to JAK2 block) are very small with the value *g* ≈ 1.73 × 10^−5^ corresponding to the minimal value (1 *− α*)*/N* in *G* related to the damping factor *α* = 0.85.

The matrix *G*_pr_ (which is exactly of rank 1) has a very simple structure with all columns being (approximately) proportional to the (local) PageRank of *G*_R_ (which is itself proportional to the global PageRank projected onto the subset of 40 nodes) and one clearly sees that the strong horizontal red lines correspond to index positions *i* of Table 1 where the corresponding index *K* is quite low below *∼* 100 corresponding to a relatively high PageRank position. The full reduced matrix *G*_R_ is numerically dominated by *G*_pr_ (but less clearly as for typical Wikipedia cases) and has at first sight a similar structure as *G*_pr_ but with somewhat smaller values. However, some of the strongest direct links (from *G*_rr_) are also visible. Similarly to the Wikipedia network of politicians as discussed in Frahm *et al*., 2016 both matrix components *G*_R_ and *G*_pr_ are not very usefully to identify the indirect links.

The indirect links are visible in the matrix *G*_qr_. As explained and shown mathematically in Frahm *et al*., 2016 they correspond to pathways where a given node *i*_1_ of the small subset points to a certain node outside the subset (in the big surrounding PPI network) which itself points eventually to another node outside the subset and comes after a finite number of iterations finally back to a different node *i*_2_ inside the subset. This provides an indirect link from *i*_1_ to *i*_2_ and the weight or strength of this indirect link is characterized by the value of the matrix element 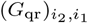. According to Figure 2 there are now also significant interactions between the two blocks of EGFR and JAK2 for the matrix *G*_qr_, sometimes with negative values (note that the matrix elements of *G*_qr_ may be negative). Figure 3 shows the the sum of the two components *G*_rr_ + *G*_qrnd_ (*G*_qrnd_ corresponds to *G*_qr_ without its diagonal elements) which confirms this observation. Actually, we consider that the elements of *G*_rr_ + *G*_qrnd_ describe best the combined direct and indirect links for the given subset.

**Fig. 3.**
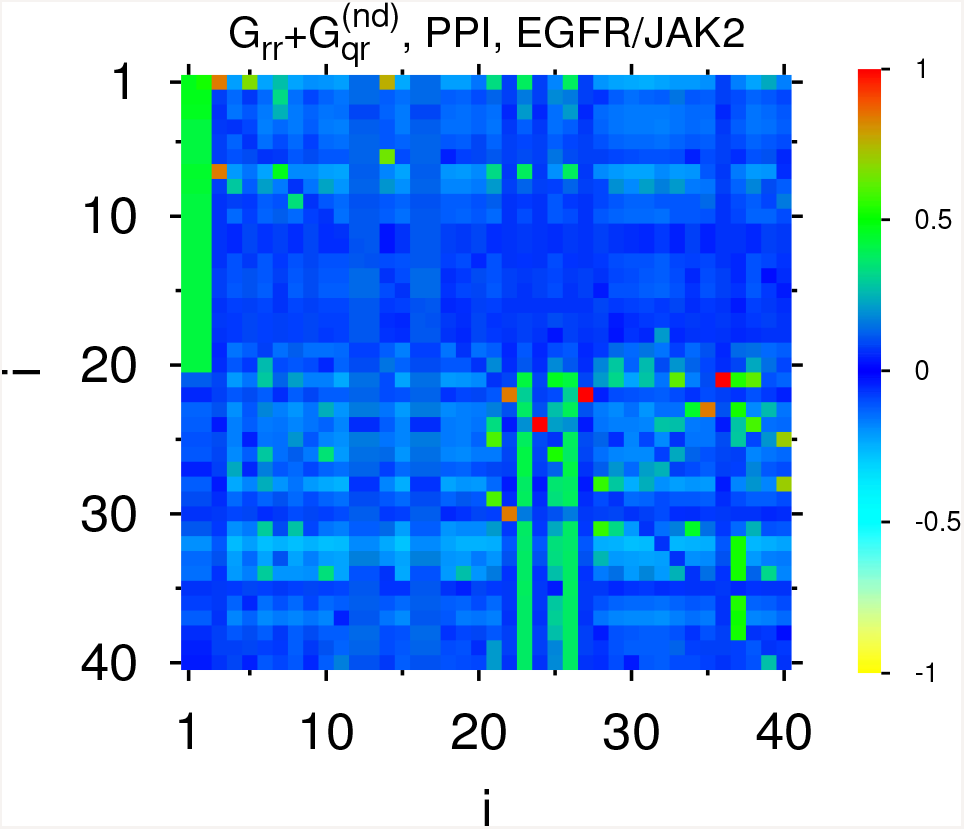
Same as in Fig. 2 but for the matrix *G*_rr_ + *G*_qrnd_, where *G*_qrnd_ is obtained from *G*_qr_ by putting its diagonal elements at zero; the weight of these two components is *W*_*rr* + *qrnd*_ = 0.227.

Due to the contribution of indirect transitions there are additional transitions between these two blocks where the four strongest additional elements of *G*_qr_ have values *g* = 0.0106 (*GRB2 P62993 (+)* to *JAK2 O60674 (+)*); *g* = 0.0099 (*GRB2 P62993 (+)* to *STAT1 P42224 (+)*); *g* = 0.0059 (*GAB1 Q13480 (+)* to *GTF2I P78347 (+)*); *g* = 0.0039 (*PIK3R1 P27986 (+)* to *GTF2I P78347 (+)*). There are also 11 additional transitions with *g* > 0.1. Thus even if the weight of *G*_qr_ is not high it provides important new indirect interactions between proteins from the EGFR block to the JAK2 block.

The situation is even more striking when we consider the transitions from the JAK2 block to the EGFR block. There are no direct links between these blocks in this direction from the global network but due to the construction of the Google matrix described above there are still numerically very small values *g* for the matrix elements of *G*_rr_ due to dangling nodes (nodes with no outgoing links) with *g* = 1*/N* ≈ 1.15 × 10^−4^ (in certain columns) or due to the damping factor term (1 *− α*)*/N ≈* 1.73 × 10^−5^ (for the other columns). On the other side concerning the indirect links described by *G*_qr_ we find rather significant transitions from the JAK2 block to the EGFR block with the four largest values: *g ≈* 0.0122 (*CCR2 P41597 (+)* to *EGFR P00533 (-)* and to *ESR1 P03372 (+)*); *g ≈* 0.006 (*CSF2RA/CSF2RB SIGNOR-C212 (+)* to *ESR1 P03372 (+)* and to *PIK3R1 P27986 (+)*). There are also 9 additional transitions with *g* > 0.001. Complete data files for the matrix elements of matrix components (for all examples) are available at Frahm and Shepelyansky, 2019c.

It is convenient to present the interactions between proteins, generated by the matrix elements of the sum of two components *G*_rr_ + *G*_qrnd_ from Figure 3, in the form of a network shown in Figure 4. To construct the network of effective friends, we select first five initial nodes which are placed on a (large) circle: the two nodes with injection and absorption (*EGFR (+)* (injection node, blue) and *JAK2 (-)* (absorption node, olive) and three other nodes with a rather top position in the *K*_*L*_ ranking: *JAK2 (+)* (related to *JAK2 (-)* with *K*_*L*_ = 1, *i* = 1, red), *GRB2 (+)* (with *K*_*L*_ = 2, *i* = 21, green) and *FES (+)* (with *K*_*L*_ = 3, *i* = 22, cyan). For each of these five initial nodes we determine four friends by the criterion of largest matrix elements (in modulus) in the same column, i.e. corresponding to the four strongest links from the initial node to the potential friends. The friend nodes found in this way are added to the network and drawn on circles of medium size around their initial node (if they do not already belong to the initial set of 5 top nodes). The links from the initial nodes to their friends are drawn as thick black arrows. For each of the newly added nodes (level 1 friends) we continue to determine the four strongest friends (level 2 friends) which are drawn on small circles and added to the network (if there are not already present from a previous level). The corresponding links from level 1 friends to level 2 friends are drawn as thin red arrows.

**Fig. 4.**
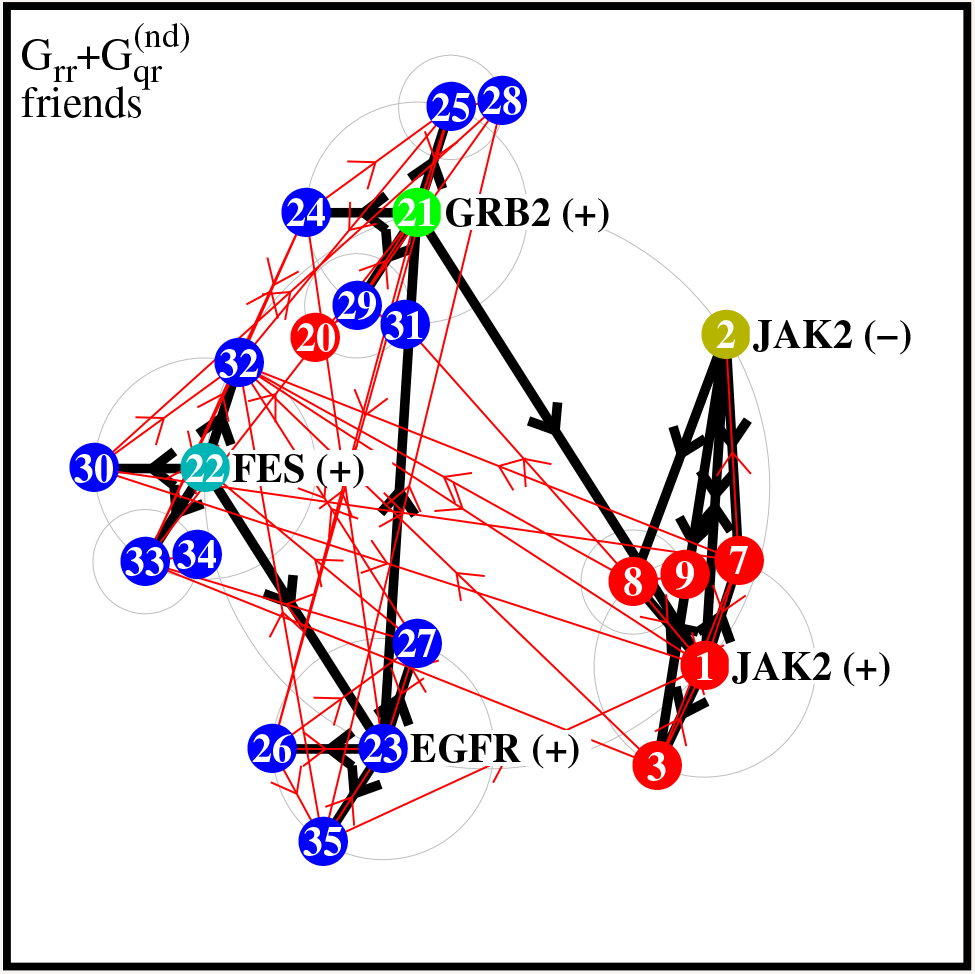
Network of friends for the subgroup of nodes given in Table 1 corresponding to injection at *EGFR P00533 (+)* and absorption at *JAK2 O60674 (-)* constructed from the matrix *G*_rr_ + *G*_qrnd_ using 4 top (friends) links per column (see text for explanations).

Each node is marked by the index *i* from the first column of Table 1. The colors of the nodes are essentially red for nodes with strong negative values of *P*_1_ (corresponding to the index *i* = 1, …, 20) and blue for nodes with strong positive values of *P*_1_ (for *i* = 21, …, 40). Only for three of the initial nodes we choose different colors which are olive for *JAK (-)*, green for *GRB2 (+)* and cyan for *FES (+)*. This procedure generates the directed friendship network shown in Figure 4.

The obtained network of Figure 4 has a rather clear separation between the two blocks related to EGFR (mainly blue nodes) and JAK2 (mainly red nodes). There is only one link of first level (black arrow) from the EGFR block (GRB2 (+)) to the JAK2 block (JAK2 (+)), Of course, there are other strong direct transitions from the EGFR block to the JAK2 block as described above, but these links are weaker than the 4 closest friends and therefore they do not appear in the network structure of Figure 4. However, we see that there are many links between the two blocks on the secondary level of red arrows.

The block of JAK2 (red nodes) is rather compact with only 6 nodes (one red node at *i* = 20 is more linked to the EGFR block). In contrast the EGFR block contains 15 (blue) nodes showing that this group of proteins is characterized by broader and more extensive interconnections. We think that such a network presentation provides a useful qualitative image of the effective interactions between the two groups of proteins.

Network figures, for this example and the other two examples discussed below, constructed in the same way using the other matrix components *G*_R_, *G*_rr_ or *G*_qr_ (instead of *G*_rr_ + *G*_qrnd_) or using strongest matrix elements in rows (instead of columns) to determine follower networks are available at Frahm and Shepelyansky, 2019c.

### 4.2 Magnetization of proteins of EGFR - JAK2 pathway

In the Ising-PPI-network each protein is described by two components which can be considered as spin up or down state. The PageRank probability of a protein is given by the sum of probabilities of its two components with *P* (*j*) = *P*_+_(*j*) + *P*_−_(*j*). It can be shown that due to the structure of the matrix transitions given by the matrices *σ*_+_, *σ*_−_, *σ*_0_ the sum of probabilities *P* (*j*) for a given protein *j* is the same as for the directed PPI network without doubling (see Supplementary Material). Thus the activation or inhibition links in the Ising-PPI-network of doubled size only redistribute PageRank probability for a given protein between up and down components. The physical meaning of these up and down component probabilities *P*_+_ and *P*_−_ is qualitatively related to the fact that on average the PageRank probability *P* of a node is proportional to the number of ingoing links. Thus *P*_+_ is proportional to the number of ingoing activation links and *P*_−_ is proportional to the number of ingoing inhibition links. Thus we can characterize each node by its normalized magnetization *M*(*j*) = (*P*_+_(*j*) − *P*_(*j*))/(*P*_+_(*j*) + *P*_(*j*)). By definition −1 ≤ *M* (*j*) ≤ 1. Big positive values of *M* mean that this protein has mainly ingoing activation links while big negative values mean that this protein has mainly inhibition ingoing links. In principle, we can also study the magnetization of CheiRank probability of proteins given by 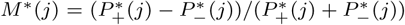 but we keep this for further investigations. We note that *M*(*j*) and *M*\*(*j*) the normalized values which are independent of the total probability *P*(*j*), *P*\*(*j*). Thus the magnetization of nodes of the reduced Google matrix remains the same as in the global network.

We take all different 38 proteins present in Table 1 and consider their magnetization (this number is smaller than 40 since for few proteins both (+) or (−) components are present in this Table). All these 38 proteins are listed in Table 2 with their local PageRank and CheiRank indices *K* and *K*\*. The distribution of these 38 proteins on the PageRank-CheiRank plane is shown in Figure 5 and the colors of the square boxes presents the values of *M*(*j*) (see caption of Figure 5). The three proteins with the strongest positive magnetizations are *PLCG1 P19174* (*M* = 0.8959), *GRB2 P62993* (*M* = 0.8899), *FES P07332* (*M* = 0.8719) and with the strongest negative values are *BCR P11274* (*M* = −0.7799), *PIK3CD O00329* (*M* = −0.7328), *PRMT5 O14744* (*M* = −0.3527). In total there are only 5 proteins of Table 2 with negative magnetization values. We attribute this to the fact that the number of inhibition links is smaller than the number of activation ones. We think that the magnetization of proteins can provide new interesting information about the functionality of proteins.

**Table 2.** Group of 38 nodes of the single protein network obtained from the group of Table 1 by removing the (+) and (−) attributes. *K* (*K\**) represent the local rank indices obtained from the PageRank (CheiRank) ordering of the single protein network. The index *i* is the same as in Table 1 where the two values *i* = 2 and *i* = 26 do not appear here since they correspond to the two nodes where both components (+) and (−) are present in Table 1.

| $K$ | $K^*$ | $i$ | Node name |
| --- | --- | --- | --- |
| 1 | 34 | 24 | PIK3CD O00329 |
| 2 | 3 | 1 | JAK2 O60674 |
| 3 | 1 | 23 | EGFR P00533 |
| 4 | 2 | 32 | PLCG1 P19174 |
| 5 | 10 | 21 | GRB2 P62993 |
| 6 | 11 | 28 | PTK2 Q05397 |
| 7 | 8 | 34 | ESR1 P03372 |
| 8 | 5 | 7 | STAT1 P42224 |
| 9 | 7 | 33 | SHC1 P29353 |
| 10 | 13 | 8 | MAP3K5 Q99683 |
| 11 | 4 | 25 | CBL P22681 |
| 12 | 25 | 31 | PIK3R1 P27986 |
| 13 | 6 | 37 | ERBB2 P04626 |
| 14 | 21 | 22 | FES P07332 |
| 15 | 12 | 5 | APOA1 P02647 |
| 16 | 9 | 19 | EZH2 Q15910 |
| 17 | 27 | 4 | ARHGEF1 Q92888 |
| 18 | 30 | 29 | GAB1 Q13480 |
| 19 | 28 | 3 | IFNGR2/INFR1 SIGNOR-C142 |
| 20 | 24 | 30 | BCR P11274 |
| 21 | 22 | 40 | CRK P46108 |
| 22 | 26 | 27 | EZR P15311 |
| 23 | 15 | 6 | CSF2RA/CSF2RB SIGNOR-C212 |
| 24 | 19 | 38 | ERBB3 P21860 |
| 25 | 33 | 36 | SHC3 Q92529 |
| 26 | 18 | 10 | CCR2 P41597 |
| 27 | 14 | 39 | NCK1 P16333 |
| 28 | 31 | 15 | ITGAL P20701 |
| 29 | 17 | 9 | STAT4 Q14765 |
| 30 | 23 | 14 | CSF2RA P15509 |
| 31 | 16 | 20 | GTF2I P78347 |
| 32 | 37 | 16 | CTLA4 P16410 |
| 33 | 36 | 13 | EPOR P19235 |
| 34 | 20 | 18 | ITGB2 P05107 |
| 35 | 38 | 17 | STAP2 Q9UGK3 |
| 36 | 32 | 11 | PRMT5 O14744 |
| 37 | 35 | 12 | STAM Q92783 |
| 38 | 29 | 35 | VAV2 P52735 |

**Fig. 5.**
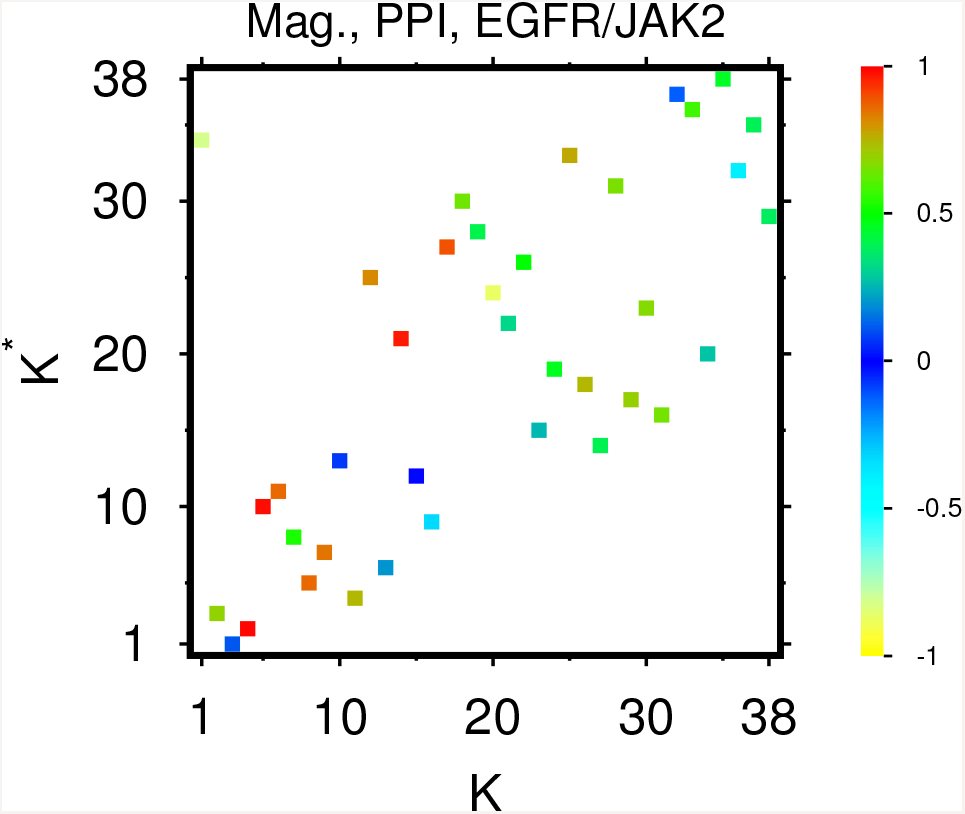
PageRank “magnetization” *M*(*j*) = (*P*_+_(*j*) − *P_*(*j*))/(*P*_+_(*j*) + *P_*(*j*)) of proteins of Table 2 shown on the PageRank-CheiRank plane (*K, K\**) of local indices; here *j* represents a protein node in the initial single protein network and *P±*(*j*) are the PageRank components of the Ising-PPI-network (see text). The values of the color bar correspond to *M/* max |*M*| with max |*M*| = 0.896 being the maximal value of |*M*(*j*)| for the shown group of proteins.

### 4.3 Examples of other protein pathways

We also consider two other proteins pairs for injection (pumping) and absorption which we analyzed in the same way. Again we compute the vector *V*_0_ = *G W*_0_ where *W*_0_ has only two non-zero components being 1 at the pumping node and −1 at the absorption node, we solve the inhomogeneous PageRank equation (1) to obtain the linear response vector *P*_1_ from which we determine a set of 40 nodes composed with 20 strongest negative and 20 strongest positive values. In order to ensure that the two initial injection and absorption nodes also belong to this subset we eventually replace the node at position 20 for strongest positive and/or negative values with the injection and/or absorption node respectively. Here we only show and discuss the list of the obtained subsets and the effective network schemes corresponding to Table 1 and Figure 4 for these two examples while Tables and Figures analogous to Table 2, Figures 1, 2, 3, 5 are given in the Supplementary Material.

First we discuss the case of injection at *MAP2K1 Q02750 (+)* and absorption at *EGFR P00533 (-)*. The protein MAP2K1 is a member of the dual-specificity protein kinase family that acts an integration point for multiple biochemical signals. There is no direct link between MAP2K1 and EGFR. The global PageRank indices of these two nodes are *K* = 84 (PageRank probability *P* (84) = 0.0009794) for *MAP2K1 Q02750 (+)* and *K* = 136 (PageRank probability *P* (136) = 0.0007817) for *EGFR P00533 (-)*. The subset of most sensitive proteins obtained from the LIRGOMAX algorithm for this protein pair is given in Table 3. These proteins are different from those of Table 1. We note now that the injection and absorption proteins have lower positions in the rank indices *K*_*L*_ and *i* of Table 3. We attribute this somehow unexpected result of the *P*_1_ ranking to rather nontrivial vortex flows on the Ising-PPI-network.

**Table 3.** Same as in Table 1 but for injection (pumping) at *MAP2K1 Q02750 (+)* and absorption at *EGFR P00533 (-)*.

| $i$ | $K_L$ | $K$ | Node name |
| --- | --- | --- | --- |
| 1 | 15 | 172 | FES P07332 (+) |
| 2 | 16 | 29 | GRB2 P62993 (+) |
| 3 | 17 | 90 | EGFR P00533 (+) |
| 4 | 18 | 3 | PIK3CD O00329 (-) |
| 5 | 19 | 126 | CBL P22681 (+) |
| 6 | 20 | 648 | EZR P15311 (+) |
| 7 | 21 | 136 | EGFR P00533 (-) |
| 8 | 22 | 38 | PTK2 Q05397 (+) |
| 9 | 23 | 30 | JAK2 O60674 (+) |
| 10 | 24 | 57 | STAT1 P42224 (+) |
| 11 | 26 | 456 | GAB1 Q13480 (+) |
| 12 | 27 | 424 | BCR P11274 (-) |
| 13 | 29 | 9 | PI3K SIGNOR-C156 (+) |
| 14 | 32 | 26 | PLCG1 P19174 (+) |
| 15 | 33 | 746 | SHC3 Q92529 (+) |
| 16 | 34 | 2398 | VAV2 P52735 (+) |
| 17 | 35 | 291 | ERBB2 P04626 (+) |
| 18 | 36 | 15 | STAT3 P40763 (+) |
| 19 | 37 | 888 | ERBB3 P21860 (+) |
| 20 | 38 | 40 | JAK1 P23458 (+) |
| 21 | 1 | 125 | CEBPA P49715 (+) |
| 22 | 2 | 144 | MAPK14 Q16539 (-) |
| 23 | 3 | 54 | GSK3B P49841 (+) |
| 24 | 4 | 543 | TAL1 P17542 (-) |
| 25 | 5 | 74 | CASP9 P55211 (-) |
| 26 | 6 | 16 | PPARG P37231 (-) |
| 27 | 7 | 1491 | ARRB2 P32121 (+) |
| 28 | 8 | 156 | MAPK3 P27361 (+) |
| 29 | 9 | 84 | MAP2K1 Q02750 (+) |
| 30 | 10 | 246 | MAPK1 P28482 (+) |
| 31 | 11 | 80 | IRS1 P35568 (-) |
| 32 | 12 | 1 | CASP3 P42574 (+) |
| 33 | 13 | 523 | KIF3A Q9Y496 (+) |
| 34 | 14 | 7 | ERK1/2 SIGNOR-PF1 (+) |
| 35 | 25 | 528 | ERG P11308 (+) |
| 36 | 28 | 106 | MEF2C Q06413 (+) |
| 37 | 30 | 826 | ANGPT2 O15123 (+) |
| 38 | 31 | 290 | TEK Q02763 (+) |
| 39 | 78 | 181 | CPT1B Q92523 (+) |
| 40 | 86 | 20 | JUN P05412 (+) |

The friendship network for this case is shown in Figure 6 (the construction method is the same as Figure 4). The 5 proteins of the initial large circle are EGFR (-) (olive), FES (+) (red), MAP2K1 (+) (cyan), MAPK14 (-) (green), CEBRPA (+) (blue). In this network we find a number of strong indirect links from the block of *MAP2K1 Q02750 (+)* (blue nodes) to *EGFR P00533 (-)* (red nodes) for which there is no direct link (e.g. from *i* = 21 to *i* = 14 proteins of Table 3). In the opposite direction from red to blue nodes there are only two strong direct matrix elements of *G*rr being from *PI3K SIGNOR-C156 (+) i* = 13 to *IRS1 P35568 (-) i* = 21 with *g* = 0.08501 and from *STAT3 P40763 (+) i* = 18 to *CASP3 P42574 (+) i* = 32 with *g* = 0.03543 with all other elements being below 1.8 × 10^−5^. However, in this direction there are 9 new indirect links with elements *g* > 0.01 and 20 with *g >* 0.005. This results in a rather dense network with many links shown in Figure 6. From the network structure we see that the proteins *i* = 25, 40 of the blue block are more closely related with proteins of the red block and inversely the proteins *i* = 10, 18, 20 of the red block are more closely related with proteins of the blue block.

**Fig. 6.**
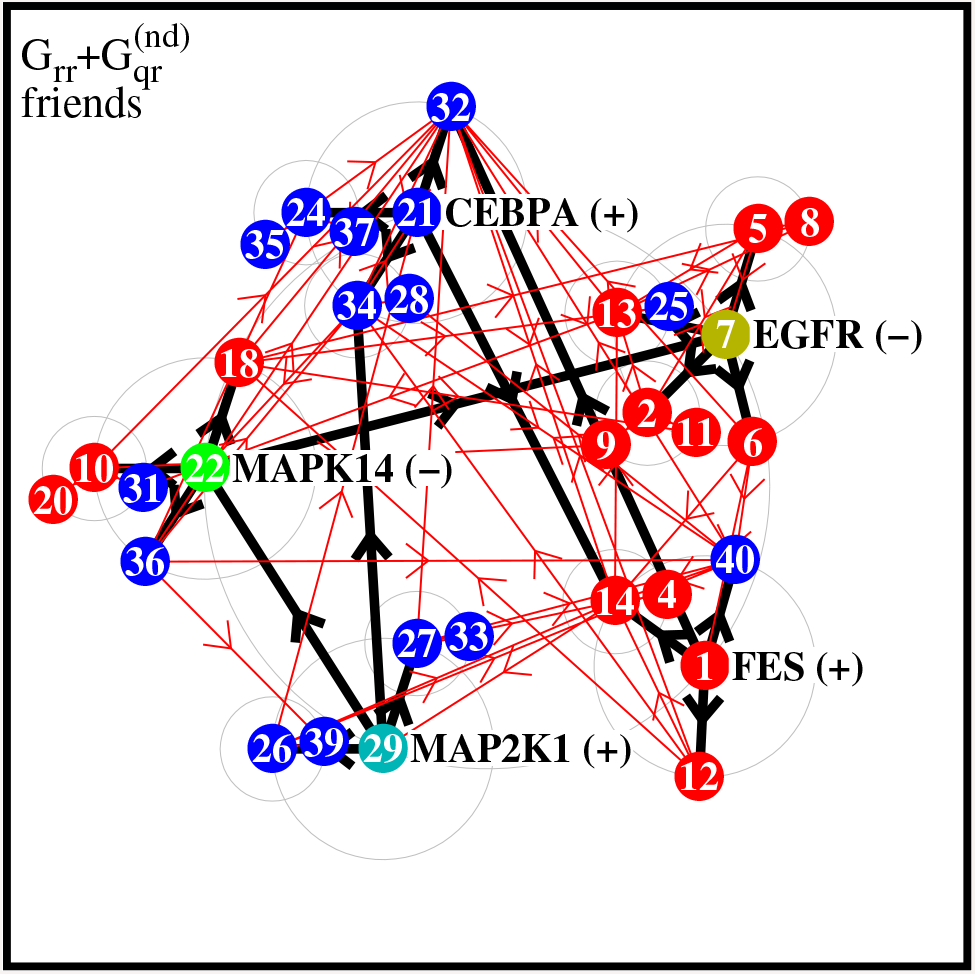
Same as Figure 4 but for the pathway of Table 3.

As a further example we also briefly discuss the pathway generated by injection at *EGFR P00533 (+)* and absorption at *PIK3CA P42336 (-)*. These two proteins are conventional bio markers of lung cancer (see e.g. Zamay *et al*., 2017). The global PageRank indices of these two nodes are *K* = 90 (PageRank probability *P* (90) = 0.0009633) for *EGFR P00533 (+)* and *K* = 1604 (PageRank probability *P* (1604) = 0.0001366) for *PIK3CA P42336 (-)*.

The most sensitive proteins obtained by the LIRGOMAX algorithm, are shown in Table 4. However, now the absorption node *PIK3CA P42336 (-)* has a very low value (in modulus) of *P*_1_ (*P*_1_ = −4.59 × 10^−5^, *K*_*L*_ = 2806) and does initially not belong to the group of nodes with 20 top strongest negative values. Therefore we replace the node *AKT3 Q9Y243 (+)* (*K*_*L*_ = 70) which was initially selected for *i* = 20 by the absorption node *PIK3CA P42336 (-)*. Furthermore, also its (+) component *PIK3CA P42336 (+)* (*P*_1_ = −0.004546 and *K*_*L*_ = 138) does not appear in Table 4 showing that the influence of *EGFR P00533 (+)* on the protein *PIK3CA P42336* is rather low.

**Table 4.**
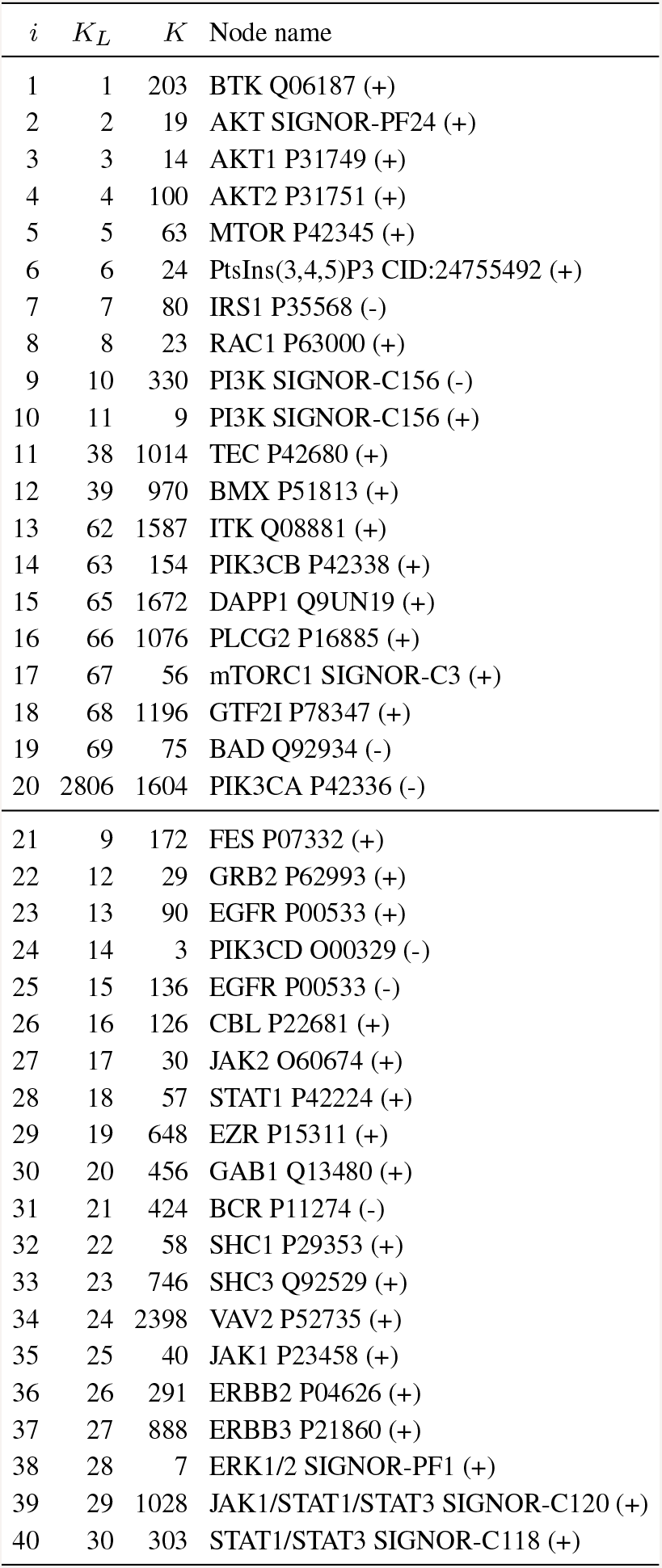
Same as in Table 1 but for injection (pumping) at *EGFR P00533 (+)* and absorption at *PIK3CA P42336 (-)*.

The friendship network structure of shown in Figure 7 shows a clear separation between the two blocks of positive (blue) and negative (red) *P*_1_ values. However, some proteins of one block happen to be closer to proteins of the other block (e.g. proteins *i* = 10, 14 from the red block are closer to the blue block and blue block protein *i* = 29 is closer to the proteins of the red block). We also note that concerning the links from the blue to the red block there are 9 significant direct transitions (matrix elements of *G*rr larger than 0.01) and 35 significant indirect *and* direct transitions (matrix elements of *G*_rr_ + *G*_qrnd_ larger than 0.01). For the opposite direction of transitions from the red to the blue block the increase is less significant but still there are new transitions due to indirect pathways (2 significant transitions for *G*_rr_ and 3 for *G*_rr_ + *G*_qr_). The significance of indirect transitions is also well visible in the friendship network of Figure 7 with many red arrows between the two blocks.

**Fig. 7.**
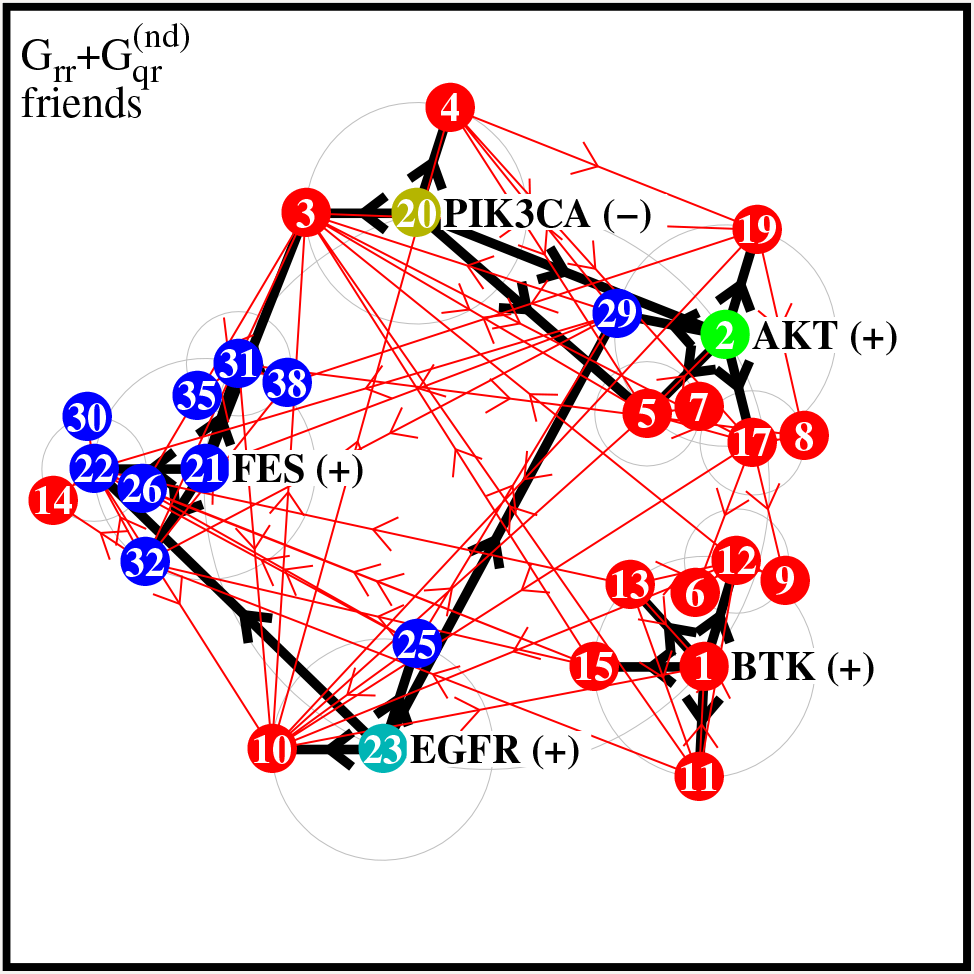
Same as Figure 4 but for the pathway of Table 4.

The same results for the original list, where the node *AKT3 Q9Y243 (+)* at position *i* = 20 has not been replaced by *PIK3CA P42336 (-)*, are available at Frahm and Shepelyansky, 2019c.

## 5 Conclusion

In this work we describe the properties of Google matrix analysis of the bi-functional SIGNOR PPI network from Perfetto, L. *et al*., 2016. The main elements of this approach are: the activation and inhibition actions of proteins on each-other are described by Ising spin matrix transitions between the protein components in the doubled size Ising-PPI-network; the recently developed LIRGOMAX (Frahm and Shepelyansky, 2019a) algorithm determines the most sensitive proteins on the pathway between two selected proteins with probability injection (pumping) at one protein and absorption at another protein; the set of most sensitive proteins are analyzed by the REGOMAX algorithm which treats efficiently all direct and indirect interactions in this subset taking into account all their effective interactions through the global PPI network. We illustrated the efficiency of this approach on several examples of two selected proteins. The obtained results show the efficiency of the LIRGOMAX and REGOMAX algorithms. We also show that the bi-functionality of protein-protein interactions leads to a certain effective magnetization of proteins which characterizes their dominant action on the global PPI network.

The executive codes and reduced Google matrix data are open and publicly available at Frahm and Shepelyansky, 2019c and interested researchers can easily study any example of a pathway between any pair of proteins from the SIGNOR network.

The described LIRGOMAX and REGOMAX algorithms can be applied also to other type of biological networks (e.g. metabolic networks discussed by Frainay *et al*., 2019).

We mention that the described Google matrix algorithms have been tested for networks with 5 million nodes and thus they can operate efficiently on other PPI networks of significantly larger size (e.g. MetaCore network which has several tens of thousands of nodes and about 2 million links). Thus we expect that the Google matrix approach, or in short Googlomics, will find broad applications for the analysis of protein-protein interactions.

## SUPPLEMENTARY MATERIAL

### S.1 Statistical properties of the SIGNOR protein-protein interactions network PPI

Using the Signor database a network of *N* = 4341 proteins with *N*_*ℓ*_ = 12547 interactions was created. In a first version, called the “single protein network”, the links do not contain the information if the interaction corresponds to activation, inhibition or is neutral/unknown. As usual we first construct an adjacency matrix with entries *A*_*ij*_ = 1 if there is a link from node *j → i* and *A*_*ij*_ = 0 if there is no such link. However, in certain rare cases there are multiple types of links between two proteins (e.g. activation and inhibition) in which case we choose *A*_*ij*_ being a multiplicity factor of 2 or 3 (instead of the usual entry 1). Once the adjacency matrix is fixed the Google matrix of this (single) protein network is constructed in the usual way: column sum normalization, taking into account the effect of dangling nodes (nodes with no outgoing link) by replacing each zero column by a uniform column with entries 1*/N* and with the application of the standard damping factor *α* = 0.85.

In Fig. S.1 we show the PageRank *P* (CheiRank *P*\*) for this single network versus the corresponding rank index *K* (*K*\*) showing a typical decay (roughly) comparable to a power law *P* ∼ 1*/K*^*β*^ (*P* * ∼ 1/(*K*\*)^*β*^) with *β ≈* 0.7 (0.8) for *K* ≥ 100 (*K* ≥* 10). Fig. S.2 shows the density of nodes in the PageRank-CheiRank plane (*K*, *K*\*) and the positions of the subgroup of nodes corresponding to Table 2 for this network.

To take into account the information about the nature of the links we use the approach of the Ising-PageRank to construct a larger network where each node is doubled with two labels (+) and (−). To construct the doubled “Ising” network of proteins each unit entry of the initial adjacency matrix is replaced by 2 × 2 matrices which are:

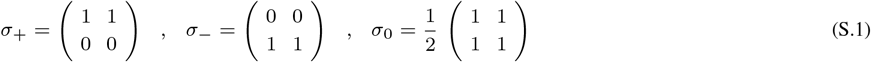

where *σ*_+_ applies to “activation”, *σ_−_* to “inhibition” and *σ*_0_ to “neutral” or “unknown”. For the rare cases with multiple types of links between two proteins we use the sum of the corresponding *σ* matrices which increases the weight of the adjacency matrix elements. After this the corresponding Google matrix is constructed in the usual way. The doubled Ising protein network corresponds to *N_I_* = 8682 nodes and *N*_*I,ℓ*_ = 27266 links (according to the non-zero entries of the used *σ* matrices).

Now the PageRank vector (of this doubles Ising network) has components *P*_+_(*j*) and *P*_−_(*j*). Due to the particular structure of the *σ* matrices (S.1) one can show analytically the exact identity *P* (*j*) = *P*_+_(*j*) + *P*_−_(*j*) where *P* (*j*) is the PageRank of the initial single protein network. For this we have to replace in Eq. (4) of (Frahm and Shepelyansky (2019)b) the value *n*_*i*_ by *n*_*ij*_ with *n*_*ij*_ = 1, 0, 1/2 for the matrix *σ*_+_, *σ*_−_ or *σ*_0_ respectively. The additional dependence of *n_ij_* on *j* takes into account that the choice of the *σ* matrix may be different for each link (and is not identical inside each row as it was the case for the model used in (Frahm and Shepelyansky (2019)b). Then the analytical argument of this work also applies in exactly the same way to the case of the doubled Ising protein network. We have also numerically verified that the identity *P* (*j*) = *P*_+_(*j*) + *P*_−_(*j*) holds up to numerical precision (*∼* 10^−13^).

As in (Frahm and Shepelyansky (2019)b) we introduce the PageRank “magnetization” by:

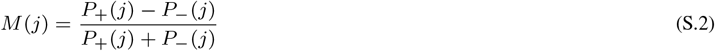

for a node *j*. The dependence of *M* (*j*) on nodes is shown in Fig. S.3 for the whole network and in Fig. 5 for the subgroup of nodes of Table 2.

**Fig. S.1.**
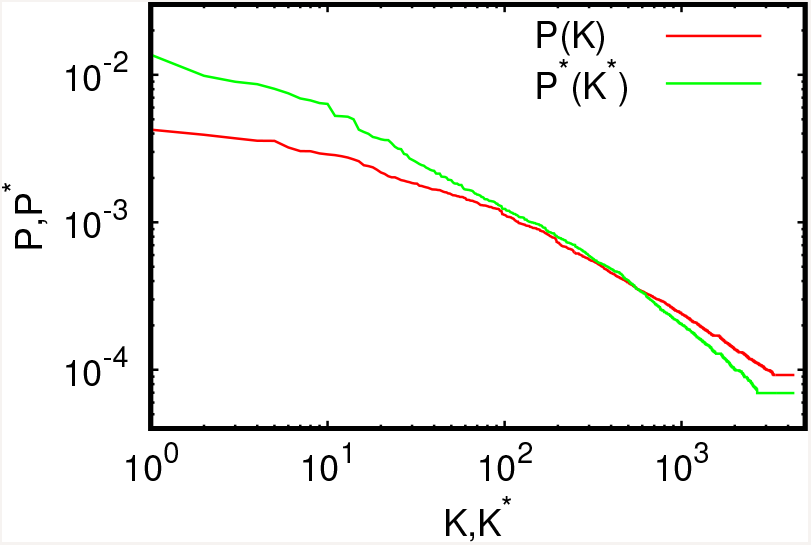
PageRank *P* (*K*) and CheiRank *P \**(*K\**) for the single protein network.

**Fig. S.2.**
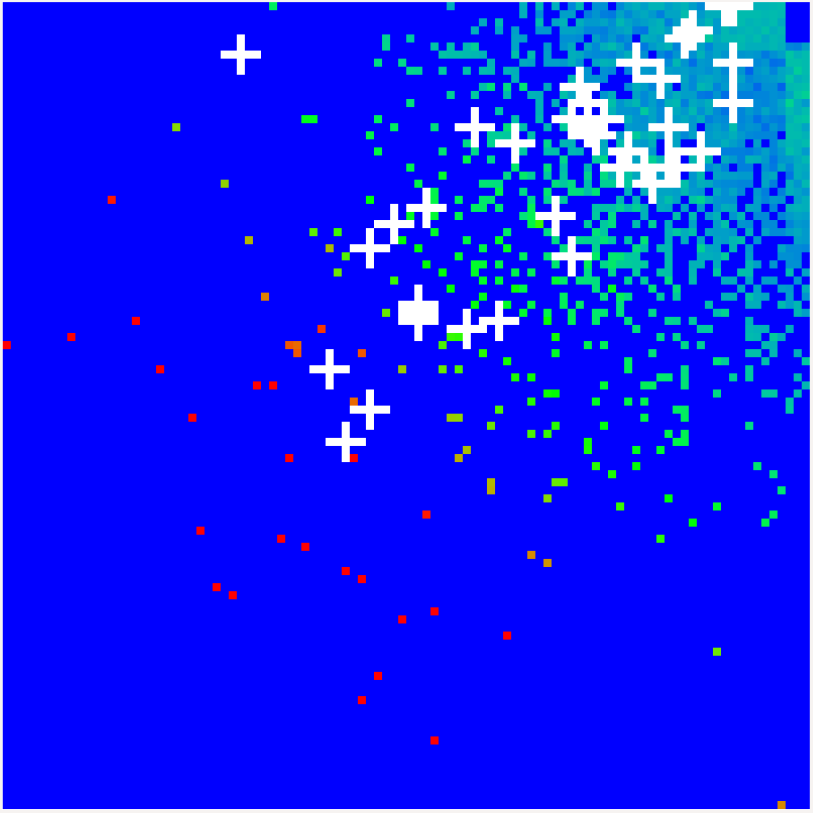
Density of nodes *W* (*K, K\**) of the single protein network on PageRank-CheiRank plane (*K, K\**) averaged over 100 × 100 logarithmically equidistant grids for 0 *≤* ln *K*, ln *K* ≤* ln *N*, the density is averaged over all nodes inside each cell of the grid, the normalization condition is *∑*_*K,K\**_ *W (K, K\**) = 1. The color bar of Fig. 2 applies (for positive values) and its values correspond to (*W/* max *W*)^1/4^. In order to increase the visibility large density values have been reduced to (saturated at) 1/16 of the actual maximum density. The *x*-axis corresponds to ln *K* and the *y*-axis to ln *K\**. The white crosses show the positions of the 38 nodes of Table 2 and in Fig. 5.

**Fig. S.3.**
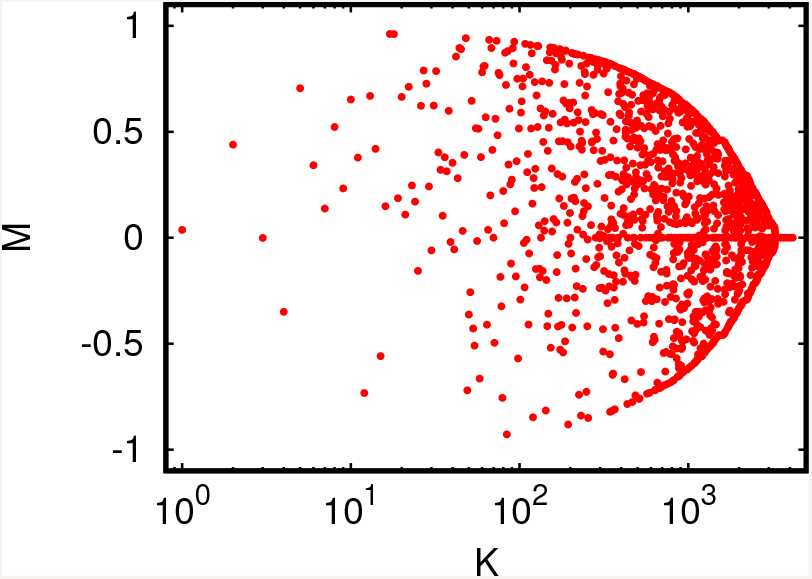
PageRank “magnetization” *M* (*j*) = (*P*_+_(*j*) − *P*_−_(*j*))/(*P*_+_(*j*) + *P*_−_(*j*)) in the Ising-PPI-network; here *j* is the node index and *K*(*j*) is the PageRank index of the initial SIGNOR network (without node doubling).

### S.2 Pathway from *MAP2K1 Q02750 (+)* **to** *EGFR P00533 (−)*

Here we present additional figures and table for this pathway discussed in subsection 4.3. Table S.1 gives the proteins (extracted from Table 3) for which the magnetization *M* is presented in Figure S.7.

**Fig. S.4.**
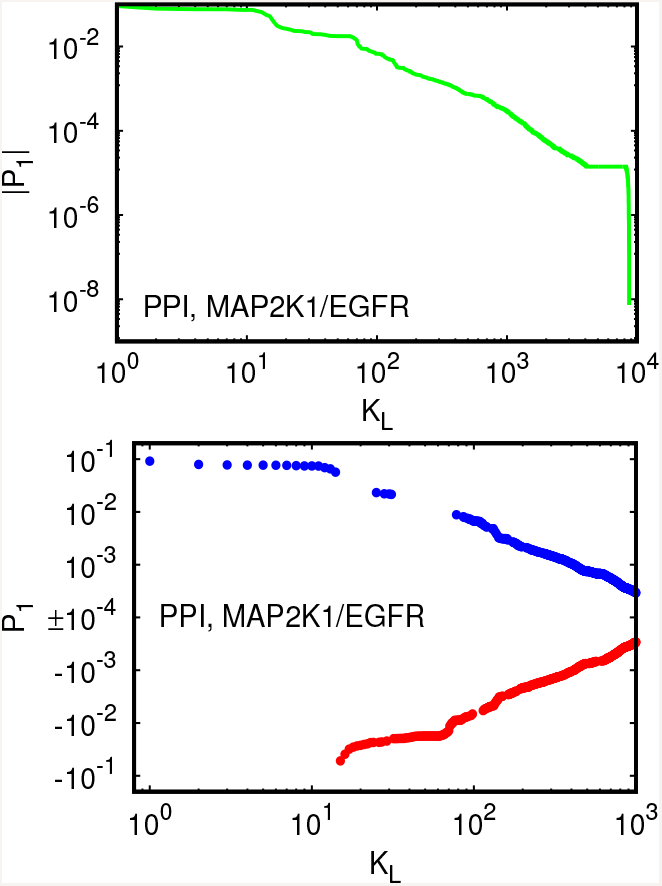
Same as in Fig. 1 but for the pathway from *MAP2K1 Q02750 (+)* to *EGFR P00533 (−)*.

**Fig. S.5.**
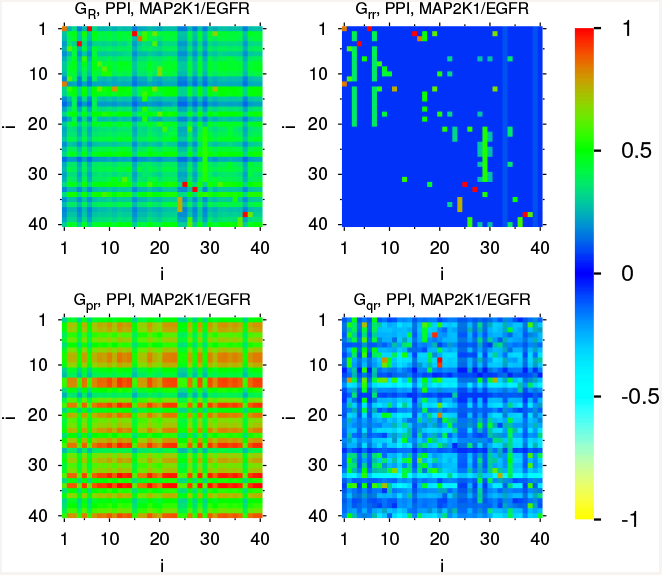
Same as in Fig. 2 but for the pathway from *MAP2K1 Q02750 (+)* to *EGFR P00533 (−)*.

**Fig. S.6.**
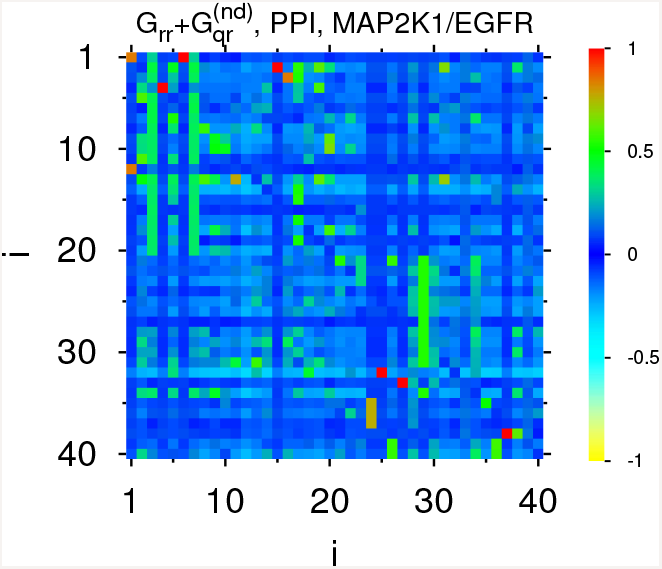
Same as in Fig. 3 but for the pathway from *MAP2K1 Q02750 (+)* to *EGFR P00533 (−)*.

**Table S.1.** Same as in Table 2 but for injection (pumping) at *MAP2K1 Q02750 (+)* and absorption at *EGFR P00533 (−)*. The index *i* is the same as in Table 3 where two values do not appear here since they correspond to the two nodes where both components (+) and (−) are present in Table 3.

| $K$ | $K^*$ | $i$ | Node name |
| --- | --- | --- | --- |
| 1 | 23 | 26 | PPARG P37231 |
| 2 | 9 | 32 | CASP3 P42574 |
| 3 | 29 | 25 | CASP9 P55211 |
| 4 | 37 | 4 | PIK3CD O00329 |
| 5 | 20 | 13 | PI3K SIGNOR-C156 |
| 6 | 8 | 18 | STAT3 P40763 |
| 7 | 5 | 34 | ERK1/2 SIGNOR-PF1 |
| 8 | 11 | 40 | JUN P05412 |
| 9 | 4 | 22 | MAPK14 Q16539 |
| 10 | 17 | 36 | MEF2C Q06413 |
| 11 | 10 | 9 | JAK2 O60674 |
| 12 | 3 | 23 | GSK3B P49841 |
| 13 | 6 | 3 | EGFR P00533 |
| 14 | 18 | 21 | CEBPA P49715 |
| 15 | 7 | 14 | PLCG1 P19174 |
| 16 | 19 | 2 | GRB2 P62993 |
| 17 | 21 | 8 | PTK2 Q05397 |
| 18 | 22 | 20 | JAK1 P23458 |
| 19 | 2 | 28 | MAPK3 P27361 |
| 20 | 31 | 31 | IRS1 P35568 |
| 21 | 14 | 10 | STAT1 P42224 |
| 22 | 1 | 30 | MAPK1 P28482 |
| 23 | 12 | 29 | MAP2K1 Q02750 |
| 24 | 13 | 5 | CBL P22681 |
| 25 | 15 | 17 | ERBB2 P04626 |
| 26 | 27 | 1 | FES P07332 |
| 27 | 39 | 39 | CPT1B Q92523 |
| 28 | 26 | 38 | TEK Q02763 |
| 29 | 25 | 24 | TAL1 P17542 |
| 30 | 33 | 11 | GAB1 Q13480 |
| 31 | 28 | 12 | BCR P11274 |
| 32 | 30 | 6 | EZR P15311 |
| 33 | 38 | 33 | KIF3A Q9Y496 |
| 34 | 16 | 35 | ERG P11308 |
| 35 | 24 | 19 | ERBB3 P21860 |
| 36 | 36 | 15 | SHC3 Q92529 |
| 37 | 35 | 37 | ANGPT2 O15123 |
| 38 | 34 | 27 | ARRB2 P32121 |
| 39 | 32 | 16 | VAV2 P52735 |

**Fig. S.7.**
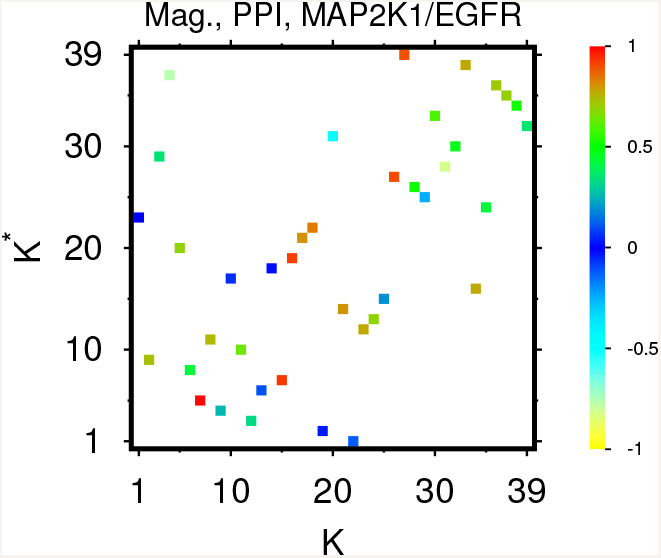
Same as in Fig. 5 but for the pathway from *MAP2K1 Q02750 (+)* to *EGFR P00533 (−)* with proteins from Table S.1; the maximal magnetization used in the color bar normalization is *M*_*max*_ = 0.961

### S.3 Pathway from *EGFR P00533 (+)* to *PIK3CA P42336 (−)*

Here we present additional figures and table for this pathway discussed in subsection 4.3. Table S.2 gives the proteins (extracted from Table 4) for which the magnetization *M* is presented in Figure S.11.

**Fig. S.8.**
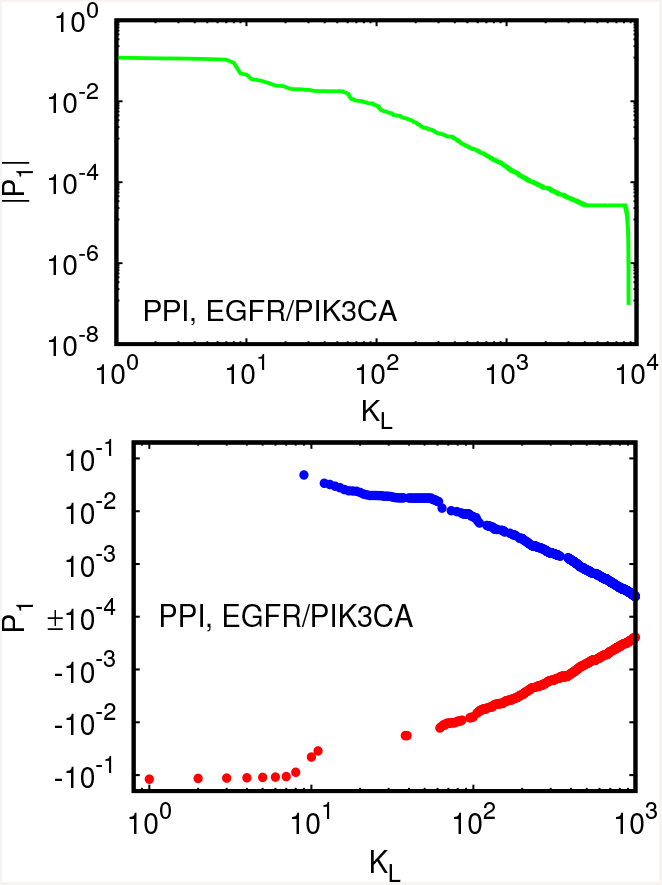
Same as in Fig. 1 but for the pathway from *EGFR P00533 (+)* to *PIK3CA P42336 (−)*.

**Fig. S.9.**
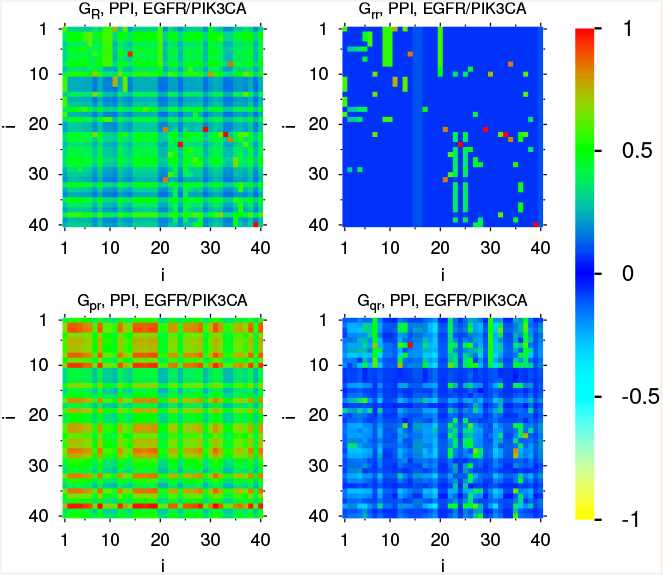
Same as in Fig. 2 but for the pathway from *EGFR P00533 (+)* to *PIK3CA P42336 (−)*.

**Fig. S.10.**
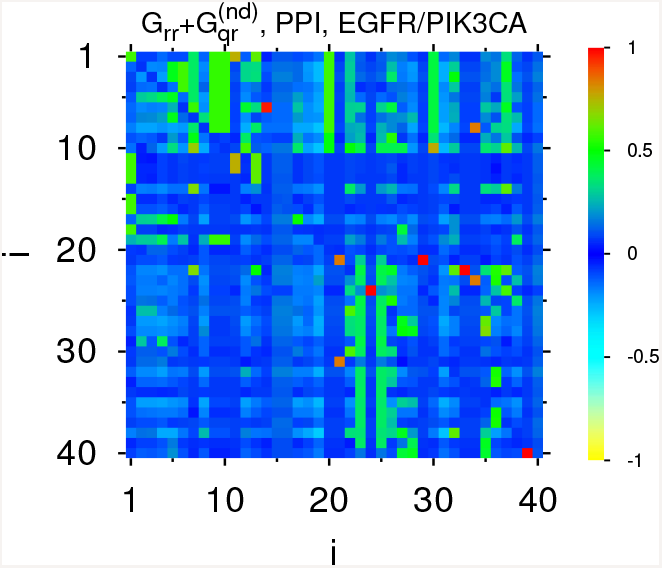
Same as in Fig. 3 but for the pathway from *EGFR P00533 (+)* to *PIK3CA P42336 (−)*.

**Table S.2.** Same as in Table 2 but for injection (pumping) at *EGFR P00533 (+)* and absorption at *PIK3CA P42336 (−)*. The index *i* is the same as in Table 4 where two values do not appear here since they correspond to the two nodes where both components (+) and (−) are present in Table 4.

| $K$ | $K^*$ | $i$ | Node name |
| --- | --- | --- | --- |
| 1 | 2 | 2 | AKT SIGNOR-PF24 |
| 2 | 1 | 3 | AKT1 P31749 |
| 3 | 35 | 24 | PIK3CD O00329 |
| 4 | 16 | 9 | PI3K SIGNOR-C156 |
| 5 | 3 | 38 | ERK1/2 SIGNOR-PF1 |
| 6 | 8 | 6 | PtsIns(3,4,5)P3 CID:24755492 |
| 7 | 14 | 17 | mTORC1 SIGNOR-C3 |
| 8 | 6 | 27 | JAK2 O60674 |
| 9 | 13 | 8 | RAC1 P63000 |
| 10 | 4 | 23 | EGFR P00533 |
| 11 | 7 | 5 | MTOR P42345 |
| 12 | 15 | 22 | GRB2 P62993 |
| 13 | 28 | 19 | BAD Q92934 |
| 14 | 25 | 14 | PIK3CB P42338 |
| 15 | 17 | 20 | PIK3CA P42336 |
| 16 | 18 | 35 | JAK1 P23458 |
| 17 | 29 | 7 | IRS1 P35568 |
| 18 | 10 | 28 | STAT1 P42224 |
| 19 | 12 | 32 | SHC1 P29353 |
| 20 | 5 | 4 | AKT2 P31751 |
| 21 | 9 | 26 | CBL P22681 |
| 22 | 11 | 36 | ERBB2 P04626 |
| 23 | 23 | 21 | FES P07332 |
| 24 | 20 | 1 | BTK Q06187 |
| 25 | 38 | 40 | STAT1/STAT3 SIGNOR-C118 |
| 26 | 31 | 30 | GAB1 Q13480 |
| 27 | 24 | 31 | BCR P11274 |
| 28 | 27 | 29 | EZR P15311 |
| 29 | 34 | 39 | JAK1/STAT1/STAT3 SIGNOR-C120 |
| 30 | 22 | 37 | ERBB3 P21860 |
| 31 | 33 | 33 | SHC3 Q92529 |
| 32 | 32 | 12 | BMX P51813 |
| 33 | 26 | 11 | TEC P42680 |
| 34 | 37 | 16 | PLCG2 P16885 |
| 35 | 21 | 18 | GTF2I P78347 |
| 36 | 19 | 13 | ITK Q08881 |
| 37 | 36 | 15 | DAPP1 Q9UN19 |
| 38 | 30 | 34 | VAV2 P52735 |

**Fig. S.11.**
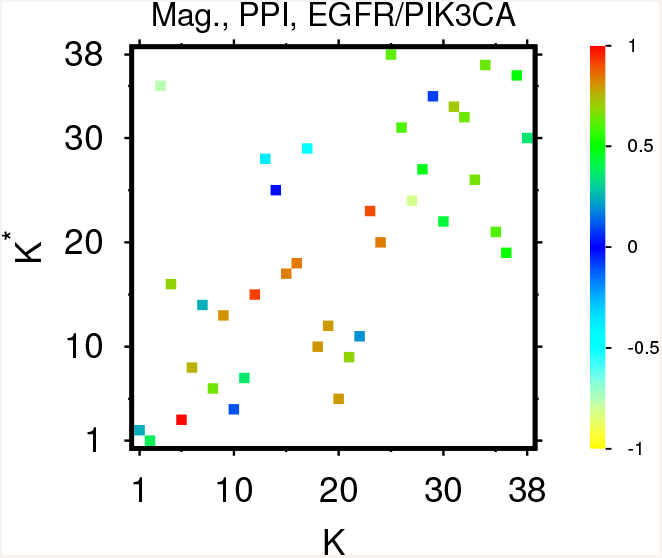
Same as in Fig. 5 but for the pathway from *EGFR P00533 (+)* to *PIK3CA P42336 (−)* with proteins from Table S.2; the maximal magnetization used in the color bar normalization is *M*_*max*_ = 0.961

